# Optimal Gene Panel Selection for Targeted Spatial Transcriptomics Experiments

**DOI:** 10.1101/2025.10.08.681071

**Authors:** Haoran Lu, Luyang Fang, Orlando Zeng, Ping Ma, Wenxuan Zhong, Guo-Cheng Yuan

## Abstract

Spatial transcriptomics analysis is a powerful approach for dissecting the structure of tissue microenvironment and uncovering the mechanism of cell-cell communications. However, existing technologies are limited by either spatial resolution or gene coverage. Most single-cell resolution technologies target only a few hundreds of pre-selected genes, whose choice plays an important role in the overall analysis. It remains a challenge to optimally design a gene panel to maximize the utility of spatial transcriptomics profiling. To fill this gap, we introduce a novel method, named ReconST, to automatically design optimal gene panels for spatial transcriptomics profiling. ReconST leverages information from existing scRNA-seq data, and identifies the optimal subset of genes by using a gated autoencoder. By using a high-coverage mouse brain MERFISH dataset as the reference benchmark, we showed that ReconST outperforms existing methods in terms of both reconstruction accuracy and spatial pattern preservation. As such, ReconST provides a useful and generally-applicable tool for optimal gene panel design, which in turn can significantly enhance the utility of spatial transcriptomics profiling in a wide range of biomedical investigations.

## INTRODUCTION

Spatial transcriptomics is an emerging and transformative technique that enables researchers to study gene expression while preserving spatial context, providing unprecedented insights into tissue organization and function [1-5]. The increasing availability of commercial platforms highlights the growing significance of spatial transcriptomics in biological studies [6, 7]. In contrast to traditional single-cell RNA sequencing (scRNA-seq) techniques, spatial transcriptomics stands out by preserving the spatial information, which is crucial for dissecting the structure of tissue microenvironment and uncovering cell-cell communication mechanisms [3, 8].

Current spatial transcriptomics technologies can broadly be categorized into two types: whole-transcriptome methods and targeted methods [9]. Whole-transcriptome spatial transcriptomics aims to measure genome-wide gene expression across tissue sections, often using techniques such as Slide-seq [10], 10x Genomics Visium [11, 12], Nanostring GeoMx Digital Spatial Profiler [13] and DBiT-seq [14], though these methods may sacrifice resolution or sensitivity [15].

In contrast, targeted spatial transcriptomics, such as MERFISH [16], seqFISH [17], seqFISH+ [18], Xenium [19], and STARMAP [20], focuses on profiling a predefined subset of genes, which is needed to achieve single-cell or subcellular resolution in spatial gene expression profiling [16, 18]. The advantage of single-cell resolution mapping is that it can significantly enhance the ability to study complex tissue architecture and cell-cell interactions [18, 21, 22]. However, its utility strongly depends on the biological information contained in the gene panel, which has to be pre-selected before spatial transcriptomics mapping. As such, an important but difficult task is to design the optimal gene panel for a biological problem at hand.

Although it is possible to manually select genes based on prior biological knowledge of cell-type specific marker genes or biological pathways, this approach has important practical limitations including subjectivity, irreproducibility, and strong dependence on prior knowledge. Recently, several methods have been developed to systematically design gene panels by leveraging information from existing single-cell RNA-seq (scRNA-seq) data (see ref. [12, 19] for review). Existing gene panel design methods vary on their selection criteria, including highly variable genes [16], marker genes associated with either annotated cell types [23] or original cell clusters [24], and the genes representative of the underlying manifold structure [25]. Despite these developments, it remains challenging to systematically design a gene panel that can optimally summarize the whole transcriptome information.

To overcome this challenge, we develop a novel method, named ReconST, for optimal gene panel design. Instead of targeting known marker genes, ReconST aims to unbiasedly select a subset of genes that optimally reconstruct the whole transcriptome information. By using a novel gated autoencoder model design, ReconST learns to compress and reconstruct gene expression profiles, ensuring that the selected genes retain as much meaningful variation as possible. ReconST offers a principled and generally-applicable solution for constructing gene panels from scRNA-seq data that are well suited for spatial transcriptomic applications. By leveraging the information from a high-coverage spatial transcriptomics dataset [26], we systematically compared the performance of ReconST and existing methods, and showed that it outperforms existing methods both for cell-type reconstruction and for recovering the underlying spatial patterns.

## RESULTS

### Overview of ReconST

ReconST is a principled data-driven method that learns to select genes directly optimized for full-transcriptome reconstruction. ReconST aims to recover as much information from the complete expression landscape as possible. At its core is a gated autoencoder architecture, a neural network designed to compress and reconstruct gene expression profiles while automatically learning which genes are most informative (Fig. 1a-c). The gating mechanism assigns learnable importance weights to each gene, and effectively selects a subset that best supports accurate reconstruction. By training on scRNA-seq data, ReconST produces a compact panel that is later validated on spatial transcriptomics datasets, ensuring both transcriptomic fidelity and spatial relevance. While neural networks have previously been used in single cell analysis, our key innovation is the gated selection layer, which dynamically assigns importance weights to each gene. Unlike combinatorial search-based methods that require exhaustive evaluation of different gene subsets, ReconST optimizes gene selection in an end-to-end fashion using stochastic gradient descent (SGD) [27]. This enables the model to efficiently learn which genes contribute most to transcriptome reconstruction, making the process highly scalable and free from heuristic selection biases. Fig. 1d shows an example that the UMAP based on the ReconST-selected panel, and when compared to the UMAP derived from the full reference panel, we see that the selected subset successfully preserves the overall cell-type clustering structure and maintains resolution across diverse populations.

**Figure 1:**
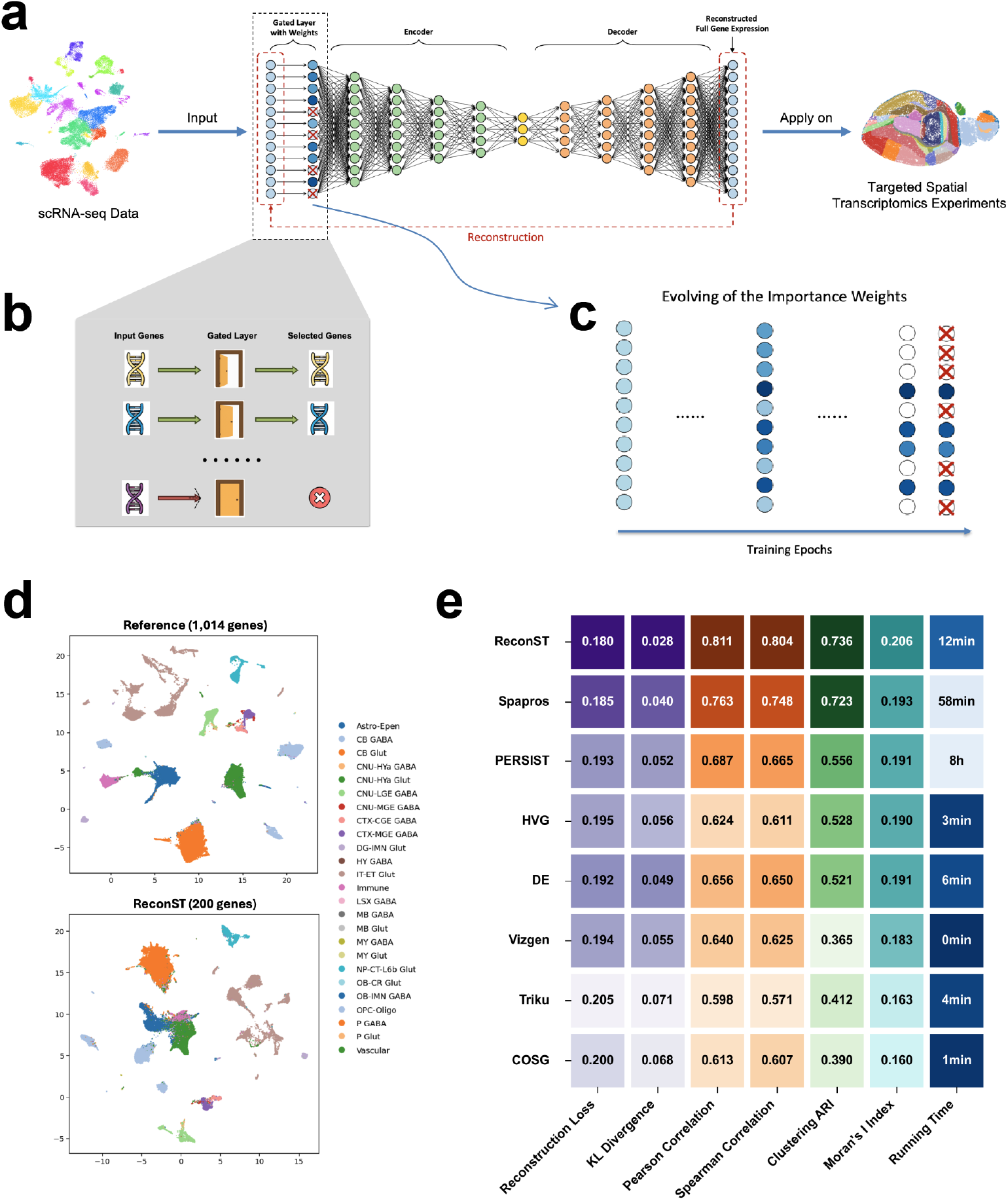
Using ReconST to design an optimal gene panel for whole mouse brain spatial transcriptomics analysis. (a) A schematic overview of the ReconST workflow using scRNA-seq data as input. (b-c) A zoomed-in view of the gated layer. (b) A schematic illustration of the gated layer; (c) An illustration of the effect of the gated layer on parameter training. Unselected genes are marked by the ‘x’ mark. (d) UMAP visualization of MERFISH data colored by annotated cell types from the reference dataset. (e) Benchmark analysis of different gene panel selection methods using the MERFISH dataset [26] as reference. Performance is evaluated by using the seven metrics.

The gated layer directly controls gene selection during training (Fig. 1b). Unlike standard autoencoders that process all input features equally, the gated layer introduces a learnable weight for every gene, allowing the model to decide which genes are essential and which can be suppressed (Fig. 1c). To further enhance efficiency and precision, we impose an L1 penalty on these weights, automatically shrinking those of non-essential genes toward zero. This sparsity-inducing regularization technique is well-established [28, 29], effectively eliminating uninformative parameters by driving their weights to exactly zero. The L1 penalty thus acts as a built-in filter, discouraging the model from assigning high importance to too many genes and ensuring that only the most informative ones are retained. After training ReconST on the scRNA-seq data, gene selection is guided by the distribution of learned weights: many are driven to zero, while non-zero weights identify genes informative for transcriptome reconstruction. These non-zero weights define the selected gene panel in a fully data-driven manner, removing the need for manual curation while preserving essential biological signals. In cases where a fixed panel size is required, the non-zero weights can be treated as importance scores, and the top-ranked genes are selected accordingly. For example, we constructed a 200-gene panel and evaluated its performance on the MERFISH dataset using reconstruction loss as the primary metric. The strength of the L1 penalty is tuned by cross-validation to balance sparsity and reconstruction accuracy [30] (see Material and Methods for details).

### Performance evaluation and benchmark using a mouse brain MERFISH dataset

To benchmark the performance of ReconST and existing methods, we utilized a publicly available dataset with a high-resolution transcriptomic and spatial atlas of the mouse brain [26]. The dataset provides both single-cell RNA sequencing (scRNA-seq) and multiplexed error-robust fluorescence in situ hybridization (MERFISH) data, allowing gene selection using scRNA-seq and validation in spatial contexts. The original scRNA-seq dataset comprises approximately 7 million cells and 19,000 genes, with 4 million high-quality cells passing stringent quality control. The data were collected from 267 donor mice, ensuring robust representation across biological replicates. While most commercially available spatial transcriptomics platforms only allow the profiling of a few hundred genes, this MERFISH dataset contains spatial profiles of 4.3 million cells measured on a panel of 1,147 genes, thereby allowing the opportunity to evaluate and compare the performance of different smaller gene panel designs in reconstruction of both transcriptomic and spatial patterns.

For fair comparison, we apply each method to select 200 genes out of the 1,147 genes used in the MERFISH dataset. To mimic real application setting, the gene panel selection was based on scRNA-seq expression data, while reserving the MERFISH data for performance evaluation. Of note, the scRNA-seq and MERFISH data were obtained from the same donor mouse, thereby providing an excellent example for performance evaluation. After preprocessing the scRNA-seq data with the SCANPY pipeline [31], a total of 31,299 cells and 3,295 genes were retained. Further analysis identified 2,019 highly variable genes, among which 1,014 genes were included in the MERFISH dataset. For model evaluation, we tested the ability of each gene panel in reconstructing the full transcriptomic and spatial information from the entire MERFISH dataset, which contains 215,278 cells and 1,014 genes after preprocessing. This carefully curated subset provides a representative foundation for evaluating our gene selection method while ensuring computational feasibility and alignment with spatial transcriptomics experiments.

We benchmarked the performance of ReconST against seven existing methods: 1. Default Panel by Vizgen [32]: The default gene panel, Vizgen 1000 Plex Gene Panel Lists for Human Brain and Mouse Brain; 2. Differentially Expressed Genes (DE) [33]: Genes selected based on differential expression analysis performed on the scRNA-seq data; 3. Highly Variable Genes (HVG): top variable genes identified from scRNA-seq data, a widely used heuristic for feature selection in single-cell studies; 4. Spapros [15]: A method that selects gene set specificity for cell type identification while capturing within-cell type expression variation; 5. PERSIST [34]: a panel design framework that integrates scRNA-seq information to preserve global transcriptomic structure, ensuring selected genes approximate the full transcriptome; 6. Triku [35]: an unsupervised method that prioritizes genes with localized expression patterns, thereby enhancing recovery of spatially informative features; and 7. COSG [36]: a correlation-based marker selection algorithm that efficiently identifies cell-type-specific genes with low redundancy. Each method was applied to select a 200 gene panel based on the curated scRNA-seq subset described above.

To ensure a comprehensive assessment of gene selection quality, we employed a series of complementary evaluation metrics. First, we computed the reconstruction loss using the mean squared error between the observed and reconstructed gene expression profiles in the MERFISH dataset. Second, we quantified the global similarity between the reconstructed and full gene expression distributions using the Kullback–Leibler (KL) divergence [37]. Third, we assessed spatial pattern preservation by calculating Moran’s I index [38], a standard measure of spatial autocorrelation, which reflects the degree to which the spatial arrangement of selected genes recapitulates that of the full transcriptome. Finally, to evaluate the cell-type resolution of each selected gene panel, we computed the Adjusted Rand Index (ARI) [39]. This was done by performing Leiden clustering [40] based on each method’s selected gene set and comparing the resulting clusters with the annotated cell-type labels. Together, these metrics provide a multifaceted evaluation of each method’s ability to recover transcriptomic structure, preserve spatial organization, and distinguish biologically meaningful cell types.

The comparative performance of the evaluated methods on the MERFISH dataset is presented in Fig. 1d. ReconST achieves the strongest overall performance with the highest Moran’s I index, indicating superior preservation of spatial autocorrelation patterns. This is followed by Spapros and PERSIST, both of which also maintain good spatial structure. ReconST additionally attains the lowest KL divergence and highest Pearson and Spearman correlations, demonstrating close agreement with the full expression distribution across both linear and monotonic relationships. Its high ARI score confirms that the selected panel retains sufficient information for accurate unsupervised cell-type discrimination.

Among the comparison methods, Spapros performs strongly as it is specifically designed to maximize cell-type specificity while preserving within-type variation, explaining its competitive ARI and spatial scores. PERSIST, which also employs an autoencoder-based approach similar to ReconST, achieves reasonable performance but requires eight hours runtime compared to ReconST’s 12 minutes. This dramatic difference stems from ReconST’s gated layer with L1 penalty that enables efficient gradient-based training, whereas PERSIST’s optimization scheme is computationally intensive. The heuristic methods HVG and DE are computationally lightweight and achieve moderate performance, but their gene selection criteria do not explicitly optimize for spatial or reconstruction objectives. The default Vizgen panel, while convenient as a pre-designed solution, shows limited performance with lower ARI and Moran’s I, likely because it was not tailored to this specific dataset’s characteristics. Triku and COSG, despite being the fastest methods, yield the weakest spatial preservation. Triku’s focus on localized expression patterns may miss genes important for global structure, while COSG’s correlation-based approach may not capture complex spatial dependencies.

Next, we compared the gene panels selected by different methods. Examination of the gene overlap patterns in Fig. 2a reveals key insights into method relationships. As expected, ReconST shows the highest overlap with Spapros (138 genes), confirming their convergence on a core set of spatially informative genes that explains their superior performance. ReconST also shares substantial overlap with DE genes (94), suggesting it effectively captures differentially expressed markers while extending beyond simple differential expression. In contrast, the poorly-performing Triku and COSG show high mutual overlap (109 genes) but limited intersection with ReconST (11 and 10 genes, respectively), indicating they miss the spatial signals that ReconST prioritizes. PERSIST, despite using an autoencoder approach like ReconST, shows only moderate overlap (108 genes), highlighting how ReconST’s gated architecture identifies a more optimal gene set. The Vizgen default panel’s moderate overlap with ReconST (43 genes) without capturing its key spatial genes underscores the advantage of ReconST’s data-driven selection over pre-designed panels. Overall, ReconST’s strong convergence with the other top performer (Spapros) and selective overlap with DE genes demonstrates its ability to identify the most spatially and biologically informative gene combinations.

**Figure 2:**
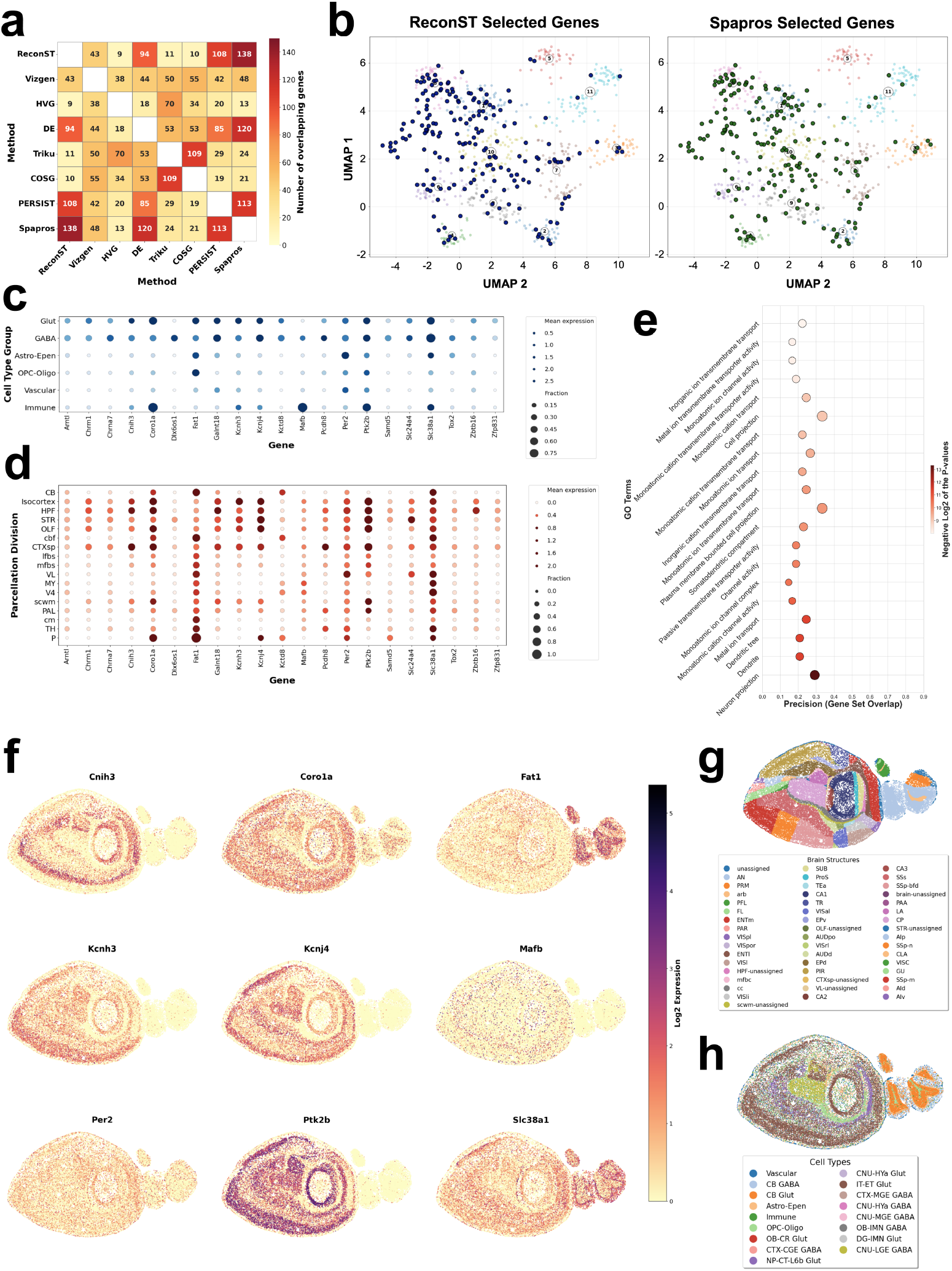
Comparison of the gene panel design by different methods. (a) Gene panel overlap of the eight methods. (b) UMAP showing expression patterns of selected genes by ReconST and Spapros, respectively. (c) Expression patterns of representative ReconST-uniquely selected genes across cell-types. (d) Expression patterns of representative ReconST-uniquely selected genes across parcellation divisions. (e) Gene Ontology enrichment analysis of the gene panel uniquely selected by ReconST. (f) Spatial patterns of representative genes. (g) Spatial patterns of different cell-types. (h) Spatial patterns of annotated anatomical parcellation divisions.

To further investigate the difference between ReconST and other methods, we compared more closely the lists of genes selected by ReconST and Spapros, which our quantitative analysis shows performs second best after ReconST. To help visualize the co-expression relationship between different genes, we calculated pairwise correlations among high-variance genes based on their spatial expression patterns, then applied UMAP dimensionality reduction to project genes into two dimensions where proximity reflects expression similarity (Fig. 2b). The distribution of genes is not random but forms several compact clusters. Within each cluster, the genes have similar expression profiles therefore contain redundant information.

Comparing to Spapros, the gene panels selected by ReconST cover more diverse patterns. Specifically, ReconST is more representative than Spapros because it covers all the major clusters, whereas the Spapros panel does not cover cluster 5 (colored red in Fig. 2b), which contains 42 genes in the full MERFISH dataset. By checking these genes in cluster 5, we find that this cluster maps to a vessel-border immune and stromal program: pan-leukocyte PTPRC/CD45, DC MIIC regulators TMEM176A/B, microglia-homeostatic SALL1, plus perivascular-fibroblast ECM markers such as COL15A1. This pattern fits border-associated macrophage/microglia engaging a perivascular fibroblast niche. Traditional panels may miss it because these cells are rare and spatially confined (with low global variance) and the genes are highly co-correlated, which redundancy-penalizing objectives down-rank.

The exclusive reach into cluster 5 motivates gene-level inspection of ReconST-unique selections. ReconST uniquely selects the two genes, Coro1a and Mafb, that reached cluster 5. Coro1a and Mafb together indicate a border myeloid program at CNS interfaces rather than a neuronal module. In Fig. 2f, Coro1a shows broad signal that aligns with cortex and major fiber tracts; biologically, CORO1A is a leukocyte actin regulator abundant in microglia, where it limits alpha-synuclein induced lysosomal stress and inflammasome activation, and it has been validated as a microglial immunohistochemical marker. Mafb appears as sparse, distributed puncta; MAFB is a macrophage-lineage transcription factor that maintains adult microglial identity and homeostasis. Interpreting these expression patterns with the cell-type and region overlays in Fig. 2g and Fig. 2h, the most parsimonious view is that both genes track microglia and border-associated macrophages in meningeal and perivascular niches, key neuroimmune hubs at brain borders. This border module can be under-prioritized by variance- or redundancy-driven panel designs because these populations are small, spatially confined, and their markers are highly correlated. By selecting these two genes, with rich spatial patterns or multi-cell-type expression, ReconST captures information that enhances the panel’s ability to represent complex cell-type relationships rather than merely recapitulating single-marker signals, reinforcing that ReconST adds biologically meaningful information beyond the other panels.

We observe that the genes uniquely selected by ReconST display rich, distributed patterns across both cell types and anatomical divisions in the mouse brain MERFISH data. For example, we observe in Fig. 2c and 2d that Per2, Kcnh3, Kcnj4, Coro1a, Slc38a1, Cnih3, Mafb exhibit clear cell-type specificity and Coro1a, Fat1, Slc38a1, Cnih3, Ptk2b, Per2, Kcnh3, exhibit clear coherent region/layer biases, indicating broad coverage captured by ReconST. The spatial expression map of these genes are shown in Fig. 2f, and they display clear spatial patterns comparing to the references in Fig. 2g and 2h.

To further investigate the biological functions of ReconST uniquely selected genes, we performed Gene Ontology (GO) enrichment analysis on them, using g:Profiler [41] via the gprofiler-official Python package, with the results shown in Fig. 2e. We see that ReconST-unique genes are strongly enriched for neuronal compartments, with the most significant terms spanning dendrite/dendritic tree, neuron projection, and cell projection. Membrane and signaling categories are also prominent, such as plasma membrane region, channel activity, passive transmembrane transporter activity, and monovalent ion transport, which are consistent with genes that shape neuronal excitability. Synaptic terms appear as well: postsynaptic membrane, neurotransmitter receptor complex, and ionotropic glutamate receptor complex, with some of these showing higher precision and larger intersections, indicating broad coverage of receptor components.

## Discussion

We have developed ReconST, a novel method for unbiased gene panel design in spatial transcriptomics analysis. ReconST uses a gated autoencoder model to select the subset of genes that optimally reconstructs whole transcriptomic information. The gated autoencoder clearly distinguishes selected from unselected genes by assigned nonzero weights to the selected genes. If the number of selected genes is pre-determined, this step can be customized by further thresholding the nonzero weights. Our benchmark analysis shows that ReconST achieves higher reconstruction accuracy, stronger spatial autocorrelation and clearer cluster resolution compared to existing methods. Further, we established a systematic benchmark that connects training on scRNA-seq data with validation on high-resolution spatial transcriptomics data (MERFISH), to evaluate how gene panel selection methods perform in realistic spatial contexts. Unlike evaluations confined to the single-cell domain, our framework measures recovery of transcriptomic content and spatial organization in actual ST data. The seven metrics (expression reconstruction, spatial autocorrelation, clustering concordance, runtime) enable fair comparison across baselines and reveal trade-offs between transcriptomic fidelity and spatial structure. This framework demonstrates the robustness of ReconST and provides a reusable pipeline for future method development and comparison.

ReconST is practical for experimental design: starting from a single-cell dataset, it outputs a compact panel ready for targeted spatial transcriptomics, reducing sequencing cost while keeping downstream analyses such as trajectory inference and differential expression reliable. By leveraging the representation learning power of deep neural networks, ReconST not only selects genes but also learns how to encode complex transcriptomic structures. Through end-to-end training, it simultaneously optimizes both feature extraction and gene selection, and the integration of data-driven gated selection with deep representation learning makes it uniquely capable of identifying the most informative and efficient gene subset for reconstruction while maintaining spatial and biological relevance. Enrichment analysis confirms that the chosen genes align with pathways that matter for the profiled tissue, and spatial maps of representative genes display coherent patterns that span several anatomical divisions. Current work is limited to one tissue and depends on a matched single-cell reference; future studies will test other samples, add multimodal information and adapt the loss for specialized tasks. Despite these constraints, ReconST offers an efficient and flexible route to information-rich panel design in spatial transcriptomics.

ReconST’s unique selections highlight biologically consequential immune programs at brain borders. Notably, Coro1a and Mafb are specifically captured and together delineate a border associated myeloid module that tracks microglia and border associated macrophages in meningeal and perivascular niches, neuroimmune hubs that are often underrepresented in variance or redundancy driven designs. Coro1a reflects leukocyte actin regulation with broad spatial signal, while Mafb marks macrophage lineage identity and microglial homeostasis, enabling sensitive readout of surveillance, cytokine exchange, and microglia associated remodeling processes. Alongside these immune markers, ReconST retains neuronal and membrane signaling genes with coherent laminar and regional biases, supporting integrated analyses of circuit level gradients and immune stromal interactions in situ. By privileging genes that carry rich spatial information rather than a fixed catalogue of markers, the panel improves detection of rare but functionally important immune states while preserving neuronal organization, thereby facilitating hypothesis generation in development, plasticity, and neurodegeneration.

## MATERIAL AND METHODS

### The ReconST Method

We present the detailed formulation of ReconST. At its core, ReconST uses an Autoencoder architecture [42], a neural network that compresses input data into a low-dimensional latent representation and reconstructs the original data from this representation. The key innovation of ReconST over traditional autoencoder is that we introduce a gate layer, positioned right after the input. The gating layer assigns non-negative learnable importance weights to each gene. Genes with zero weight are excluded from the reconstruction process, while those with higher weights are prioritized, allowing the model to focus on the most informative genes and effectively filter out noise.

Following the gate layer, the autoencoder is composed of two parts: an encoder and a decoder. The input to the encoder is the gated full gene expression profile. The encoder consists of fully connected layers with gradually decreasing dimensions to map the high-dimensional gene input into a compressed representation in a lower-dimensional latent space. Then, the decoder with increasing number of nodes then reconstructs the original input from this representation. Together, the encoder and decoder form a bottleneck architecture, in which the latent space serves as a constraint that forces the network to capture the most salient features of the data. This structure prevents the model from learning a trivial identity mapping and encourages the extraction of compact and meaningful representations. Finally, we have the output of decoder to be the reconstructed full gene expression.

Mathematically, the input gene expression vector for the *i*-th cell is denoted as ***X***_*i*_ ∈ *R*^*g*^, where *g* is the total number of genes. The key trainable gating layer is introduced immediately after the input layer, assigning a learnable scalar weight *w*_*j*_ to each gene *j*. These weights form a gating vector ***w*** = (*w*_1_, *w*_2_, ⋯, *w*_*G*_)^*T*^, which modulate the input gene expression vector ***X***_*i*_ is through element-wise multiplication to produce a gated input 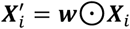, where ⊙ denotes element-wise multiplication. Genes with near-zero weights *w*_*j*_ are suppressed, while those with non-zero weights are prioritized for reconstruction. The encoder then maps the gated input ***w*** ⊙ ***X***_*i*_ to a compressed latent representation ***h***_*i*_ = *f*_*θ*_(***w*** ⊙ ***X***_*i*_) using a multi-layer perceptron structure [43]. This encoder comprises *H* sequential nonlinear transformations:

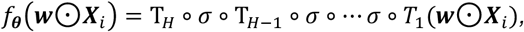

where each *T*_*j*_ represents an affine transformation *T*_*j*_(***u***) = ***v***^(*j*)^***u*** + ***b***^(*j*)^, *j* = 1, ⋯, *H* parameterized by weights 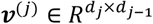 and biases 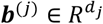. Here, *d*_*j*_ denotes the dimension of the *j*-th layer, and *σ*(⋅) is the LeakyReLU activation function [43]. The encoder parameters ***θ*** are the collection of all ***v***^(*j*)^ and ***b***^(*j*)^. Similarly, the decoder *g*_***ϕ***_, parameterized by *ϕ*, reconstruct the input from the latent representation ***h***_*i*_:

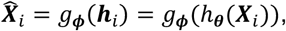

aiming to minimize the reconstruction error between the original input ***X***_*i*_ and the reconstructed output 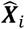.

To encourage sparsity in gene selection, we imposed an *L*_1_ penalty on the gating weights ***w***. The model is trained by minimizing a composite loss function:

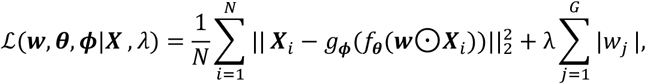

where *N* is the total number of samples, *λ* controls the strength of the *L*_1_ regularization, and ***X*** = (***X***_1_, …, ***X***_*N*_)′ denotes the whole training dataset. The first term in the loss function quantifies the reconstruction error using mean squared error, while the second term encourages sparsity by *L*_1_ regularization of the gating weights.

By optimizing this loss, ReconST learns to retain only genes with non-zero *w*_*j*_ values, which are most critical for reconstructing the full transcriptome, while discarding uninformative genes. The L1 regularization leads the weights of noninformative genes to shrink towards zero [28, 44, 45], and the selected gene subset to be sparse, reducing redundancy to achieve our goal of gene subset selection. As a result, ReconST automatically adjusts these weights to suppress irrelevant genes and emphasize those critical for accurate reconstruction of the full gene expression profile.

After model training, genes with non-zero gate weights are selected as the informative gene panel. This approach allows the model to automatically determine a data-driven subset of genes that contribute meaningfully to reconstruction. In cases where a pre-specified number of genes is required and this number is smaller than the total number of non-zero weights, the gate weights are treated as importance scores, and the top-ranking genes are selected accordingly.

### Implementation Details

#### ReconST method

We use a symmetric autoencoder structure where the size of input and gated layers both equal to the dimension of genes. This is followed by layers with 512, 512, 256, 256, and 128 nodes in the encoder. The decoder mirrors this structure with layers of 256, 256, 512, 512, and finally the same number of nodes as the gene dimension in the output layer. In our example, the gene dimension is 1,147. Each hidden layer (other than the gated layer and the output layer) is followed by a LeakyReLU [46] activation function with a negative slope of 0.3 and a dropout layer [47] with a dropout probability of 0.05. The model was trained using the Adam optimizer [48] with a learning rate of 0.001 and weight decay of 0.00001, and training proceeded for 1,000 epochs. All hyperparameters, including the learning rate, weight decay, dropout rate, and LeakyReLU slope, were selected via grid search based on a validation set separated from the training set. The *L*_1_ penalty was applied only to the training procedure, while test loss was computed using only the reconstruction term to assess the reconstruction performance.

#### Data source

We used matched single-cell RNA-seq and MERFISH resources from the Allen Brain Cell Atlas as described in [26]. The single-cell reference was the cortex subset WMB-10Xv2-CTXsp-log2.h5ad. The spatial validation set was Zhuang-ABCA-4-log2.h5ad with the accompanying cell metadata, gene table, and CCF section coordinates.

#### Preprocessing

For scRNA-seq, we removed cells with fewer than 200 detected genes and genes present in fewer than 100 cells, followed by library-size normalization to a target of 10,000 counts, log transformation, and scaling with a maximum absolute value of 10. MERFISH values were used as provided in log scale. Gene symbols were harmonized between modalities and intersected to form a common gene set, and CCF section x and y coordinates were linked to cell identifiers for spatial analyses.

#### Panel selection and compared methods

All gene panels were derived exclusively from the scRNA-seq reference, with panel size K set to 200 unless stated otherwise. ReconST is a gated autoencoder trained on scRNA-seq that assigns a non-negative gate to each gene with an L1 penalty for sparsity. The encoder comprises 512, 512, 256, 256, and 128 units with LeakyReLU slope 0.3 and dropout 0.05, and the decoder mirrors this structure. Training minimized mean-squared reconstruction error plus the sparsity penalty using Adam with learning rate 1×10^−3^, weight decay 1×10^−5^, batch size 256, and an 80:20 train-validation split; the panel consists of the top-K gated genes. Baselines derived from scRNA-seq included highly variable genes using SCANPY, differential-expression markers identified by Wilcoxon after Leiden clustering and aggregated to 200 unique genes, Triku after PCA with cosine neighbors, COSG with group definitions from Leiden, Spapros with a random-forest selector using provided cell-type labels, and PERSIST trained with a hurdle loss on binarized and log-CPM layers. The Vizgen-1k list served as a fixed vendor baseline after intersecting with MERFISH genes and subsampling to K when necessary. A random-gene panel was used only for preliminary checks and is not reported.

#### Evaluation

All quantitative evaluation was performed on MERFISH only. For each method’s panel we masked unselected genes in MERFISH and used a shared decoder to reconstruct the full common-gene expression, providing a uniform “given K genes, how much of the transcriptome can be recovered” comparison. We reported mean-squared error across cells and genes, gene-wise Kullback–Leibler divergence between epsilon-smoothed cell-wise distributions averaged across genes, and gene-wise Pearson and Spearman correlations averaged across genes. Spatial preservation was assessed with Moran’s I computed using Squidpy and the CCF section coordinates, averaged across genes. Cell-type separability was assessed by running PCA, neighbors, and Leiden on each panel’s genes and computing the Adjusted Rand Index against the provided taxonomy. Runtimes reflect the wall-clock time required to obtain each panel on scRNA-seq.

#### Software and code availability

Analyses were performed in Python. Scanpy handled preprocessing, highly variable genes, differential expression, neighbors, and Leiden. Squidpy was used for Moran’s I. Scikit-learn provided evaluation metrics including the Adjusted Rand Index. PyTorch was used for the autoencoder and reconstruction loss, and NumPy, SciPy, and pandas supported numerical operations and data handling; matplotlib and seaborn were used for plotting. Spapros, COSG, Triku, and PERSIST were run from their public Python implementations. Code for the ReconST method will be publicly available on GitHub at https://github.com/haoranlustat/ReconST), and will later be made available as a open Python package.

